# Heterogeneity in Neutrophil Extracellular Traps from Healthy Human Subjects

**DOI:** 10.1101/2023.11.03.565547

**Authors:** Margaret S. Collins, Michelle A. Imbrogno, Elizabeth J. Kopras, James A. Howard, Nanhua Zhang, Elizabeth L. Kramer, Kristin M. Hudock

**Affiliations:** Division of Pulmonary, Critical Care & Sleep Medicine, Department of Medicine, University of Cincinnati College of Medicine, Cincinnati, OH USA; Department of Pharmacology & Systems Physiology, University of Cincinnati, Cincinnati, OH, USA; Department of Pediatrics, University of Cincinnati College of Medicine, Cincinnati, OH USA; Division of Biostatistics and Epidemiology, Cincinnati Children’s Hospital Medical Center, Cincinnati, OH, United States; Division of Pediatric Pulmonary Medicine, Cincinnati Children’s Hospital Medical Center, Cincinnati, OH USA; Division of Pulmonary Biology, Cincinnati Children’s Hospital Medical Center, Cincinnati, OH USA

**Keywords:** NET heterogeneity, neutrophil extracellular traps, DNA, neutrophil elastase

## Abstract

Neutrophil Extracellular Traps (NETs), a key component of early defense against microbial infection, are also associated with tissue injury. NET composition has been reported to vary with some disease states, but the composition and variability of NETs across many healthy subjects provides a critical comparison that has not been well investigated. We evaluated NETs from twelve healthy subjects of varying ages isolated from multiple blood draws over a three and one half-year period to delineate the variability in extracellular DNA, protein, enzymatic activities, and susceptibility to protease inhibitors. We calculated correlations for NET constituents and loss of human bronchial epithelial barrier integrity, measured by transepithelial electrical resistance, after NET exposure. We found that although there was some variability within the same subject over time, the mean numbers of neutrophils, protein, LDH, serine protease activities, and cytokines IL-8, IL-1RA, and G-CSF in isolated NETs were consistent across subjects. Total DNA and double stranded DNA content in NETs were different across donors. NETs had little or no TNF*α*, IL-17A, or GM-CSF. NET DNA concentration correlated with increased NET neutrophil elastase activity and higher NET IL-1RA concentrations. NET serine protease activity varied considerably within the same donor from day-to-day. Mean response to protease inhibitors was significantly different across donors. NET DNA concentration correlated best with reductions in barrier integrity of human bronchial epithelia. Defining NET concentration by DNA content correlates with other NET components and reductions in NET-driven epithelial barrier dysfunction, suggesting DNA is a reasonable surrogate measurement for these complex structures in healthy subjects.

## 1. Introduction

Neutrophils provide a first line of defense against infection (1). Polymorphonuclear leukocytes (PMN) are actively recruited to the site of injury or infection by chemoattractants produced by the host and by invading microbes (2, 3). Stimulated PMN responses include phagocytosis, release of oxidative burst, and release of antimicrobial peptides into the cytoplasm (4). Histone arginine and methylarginine are converted to citrulline leading to structural changes in the neutrophil chromatin (5, 6). Subsequently, decondensation of nuclear contents and expansion of the nuclei leads to an explosive extracellular release of neutrophil extracellular traps (NETs) via NETosis (7).

NETs are made up of DNA strands studded with bioactive compounds such as proteases (8). Serine proteases neutrophil elastase (NE), proteinase 3 (PR3) and cathepsin G (CG) are present in NETs and can be bactericidal (9). The cytokine IL-8 is another immunomodulatory protein present in NETs (10). While the relative abundance of the protein components has been reported, little is known about the variability in NET composition and enzymatic activities across healthy human NET donors (11). Likewise, the degree to which these components correlate with NET-induced lung tissue injury have not been investigated. We previously reported that exposing primary normal human bronchial epithelial cells (HBE) to NETs led to a breakdown in epithelial barrier integrity including decreases in transepithelial electrical resistance (TEER) (10). NET exposure also drove epithelial secretion of select inflammatory cytokines, TNF-*α* and IL-8, via activation of the IL-1 pathway (12). In this study, we sought to characterize isolated NETs from normal human donors, including analysis of DNA and protein content, enzymatic activities, and susceptibility to selective inhibitors. We then correlated these NET components with the loss of barrier integrity in NET-exposed HBE.

## 2. Results

### 2.1. Mean neutrophil numbers are consistent across subjects

We freshly collected peripheral blood from healthy male and female subjects over 3 1/2 years, from 2020-2023. Donor characteristics and the number of blood collections for each donor are described in Supplemental Tables 1 & 2. Negative bead selection was used to obtain highly enriched populations of neutrophils with a mean cell viability of 94.1%. In our hands, negative bead selection, though more costly, resulted in a homogenous cell population with less spontaneous activation of neutrophils compared to density gradient separation (Supplemental Figure 1 A-B). Immediately after isolation, neutrophils were exposed to Phorbol 12-myristate 13-acetate (PMA) and NETs were isolated. To determine which NET components are likely causing lung tissue injury, we sought to delineate the extent to which NET content varies across different healthy donors and within the same donor on different days. Isolated human NETs were assessed for DNA content, protein concentration, lactate dehydrogenase (LDH), serine proteases (NE, CG, and PR3), myeloperoxidase (MPO), and cytokines. In Figure 1A, neutrophils per milliliter peripheral blood from healthy donors was compared. Neutrophil concentrations ranged from 9.4e5 to 5e6/ml and the mean number of neutrophils isolated was consistent across donors. There was no significant difference in variability across donors (p=0.060).

**Figure 1.**
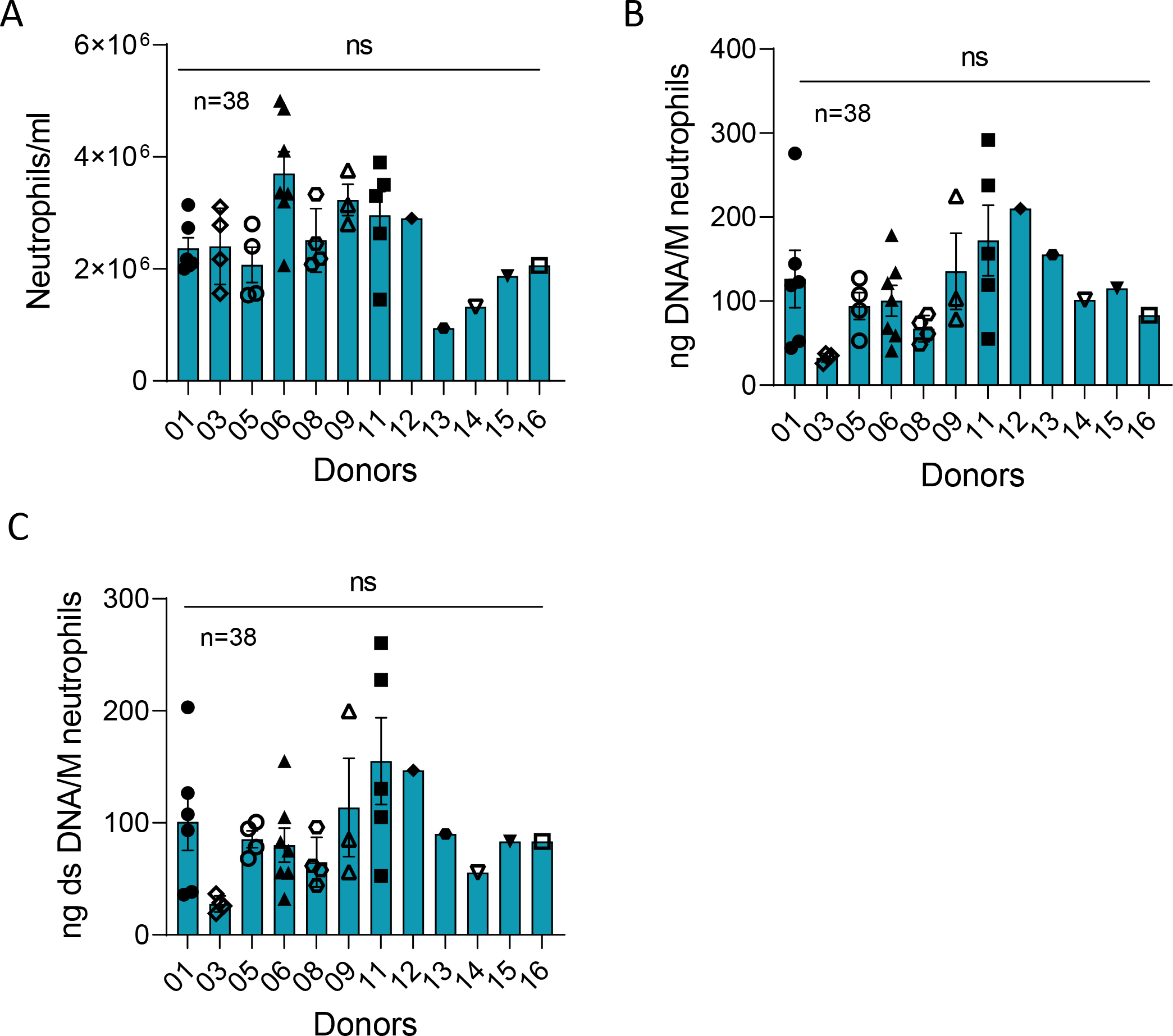
Total extracellular DNA content of NETs is similar across donors. **A)** Mean number of neutrophils recovered per ml blood drawn from 12 healthy donors in 38 collections using negative bead selection was not significantly different between donors (p=0.060). **B)** Mean NET DNA and **C)** dsDNA concentrations expressed as ng DNA/million neutrophils was not significantly different between donors (p=0.096 and 0.156, respectively). Data analyzed across subjects by one-way ANOVA.

### 2.2. DNA concentration in NETs is not significantly variable across donors

DNA serves as the backbone of NETs. Total extracellular DNA and double-stranded DNA (dsDNA) measurements of each NET preparation were averaged and expressed as ng DNA per 10^6^ neutrophils (Fig 1B). Average DNA content ranged from 25.9 to 291.9 ng/million neutrophils with a mean NET DNA of 109.73+/-182.17ng/ml. The mean NET DNA content was not significantly different across subjects (p=0.096). Four of seven donors exhibited variability from day to day in DNA suggesting that some subjects’ neutrophils may vary in their capacity to form NETs.

Interestingly, dsDNA represents an average of 29.6+/-30.32% of the total DNA found in NETs generated from healthy humans (Figure 1B-C). dsDNA content ranged from 19.2+/-260.4ng/million cells and the mean dsDNA was not significantly different across donors. Variability in dsDNA content was not significant between subjects (p=0.156).

### 2.3. NET DNA serves as a scaffold for bioactive molecules

The NET structure carries a payload of antimicrobial proteins. Total protein content of isolated NETs was measured and normalized to μg DNA of each sample (Figure 2A). Total protein in the NETs was not significantly different between donors (p=0.586), but there was variability within the same subject from day-to-day for a few donors (06, 09, 11), whereas other donors had less variability.

**Figure 2.**
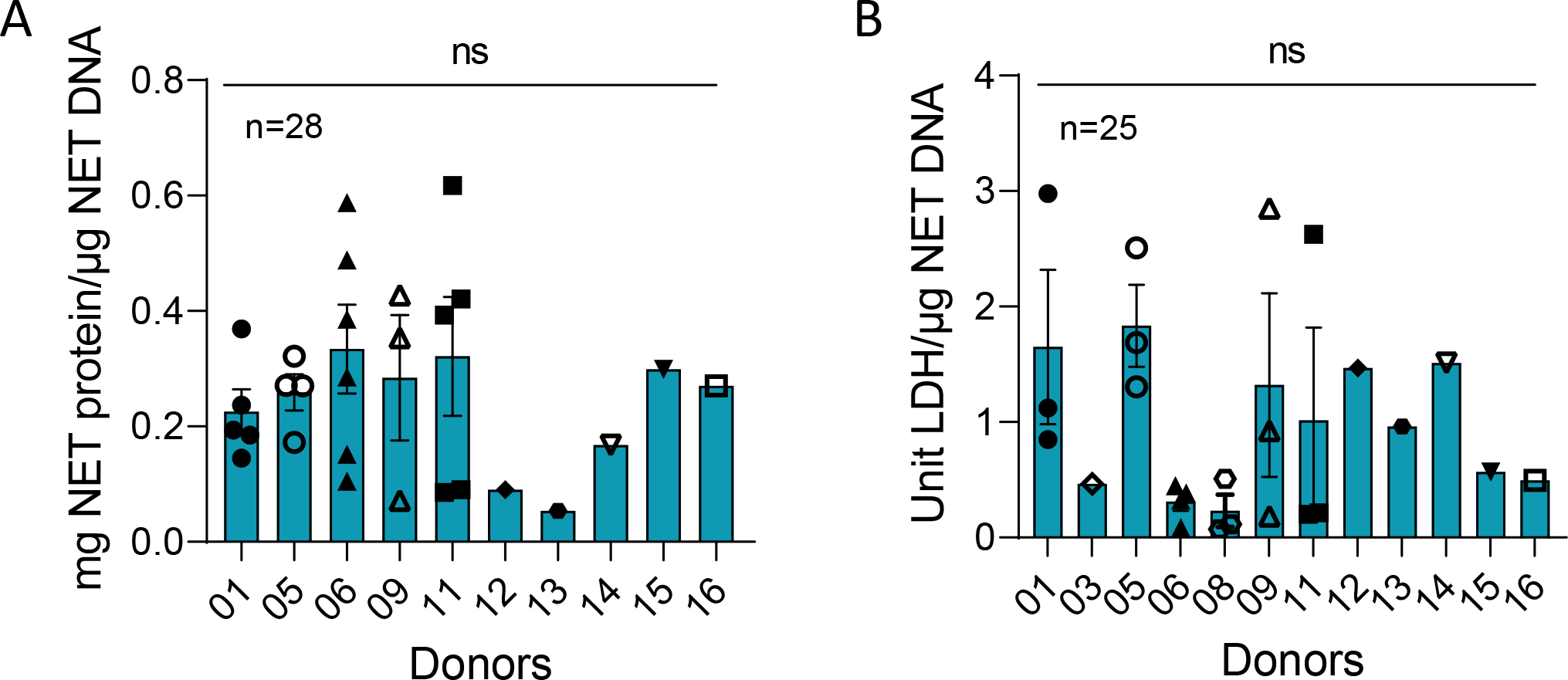
Total protein and LDH is consistent across human NETs. Mean values of protein and LDH were compared across 25 and 28 blood draws from 10 and 12 donors, respectively. Samples were normalized to μg NET DNA. **A)** No significant difference was found in protein concentration between donors (p=0.581). Data analyzed across subjects by one-way ANOVA. **B)** LDH levels, also normalized to NET DNA, were not significantly different across donors (p=0.817). Data analyzed across subjects by Kruskal Wallis.

LDH was consistently present in our isolated NETs produced by stimulation of neutrophils with 25nM PMA. LDH is released into the supernatant in response to cell membrane damage and is an indicator of cytotoxicity (17). There was no significant difference in the mean (p=0.817) of LDH across individuals (Figure 2B).

NE, CG, and PR3, stored in the azurophilic granules of PMNs, are structurally related, have antibacterial properties, and are critical to the destruction of pathogens (1). These bioactive compounds are part of the milieu of proteins associated with NETotic DNA. In figures 3A-D, the greatest serine protease activity was contributed by NE with a mean of 35.04+/-81.6mU/μg NET DNA. CG and PR3 concentrations were 1.56+/- 6.99μU/μg and 4.4+/-20.5μU NET DNA, respectively. The mean NE, CG and PR3 activities were not significantly different across donors. Although there was no statistically significant difference in variability across donors for NE (p=0.129), CG (p=0.332) PR3 (0.717) or MPO (p=0.143) activities, there was variability within repeated measures from the same NET donor (NETs isolated on different days). MPO is also stored in azurophilic granules, is antimicrobial, can be a source of inflammation, and be involved in NET formation (18, 19). Mean MPO activity measured 7.99+/-1.63e-5pM/min/ml per μg NET DNA. In correlations of serine protease and MPO activities to NET DNA concentration, only NE was positively correlated (Figure 5A).

**Figure 3.**
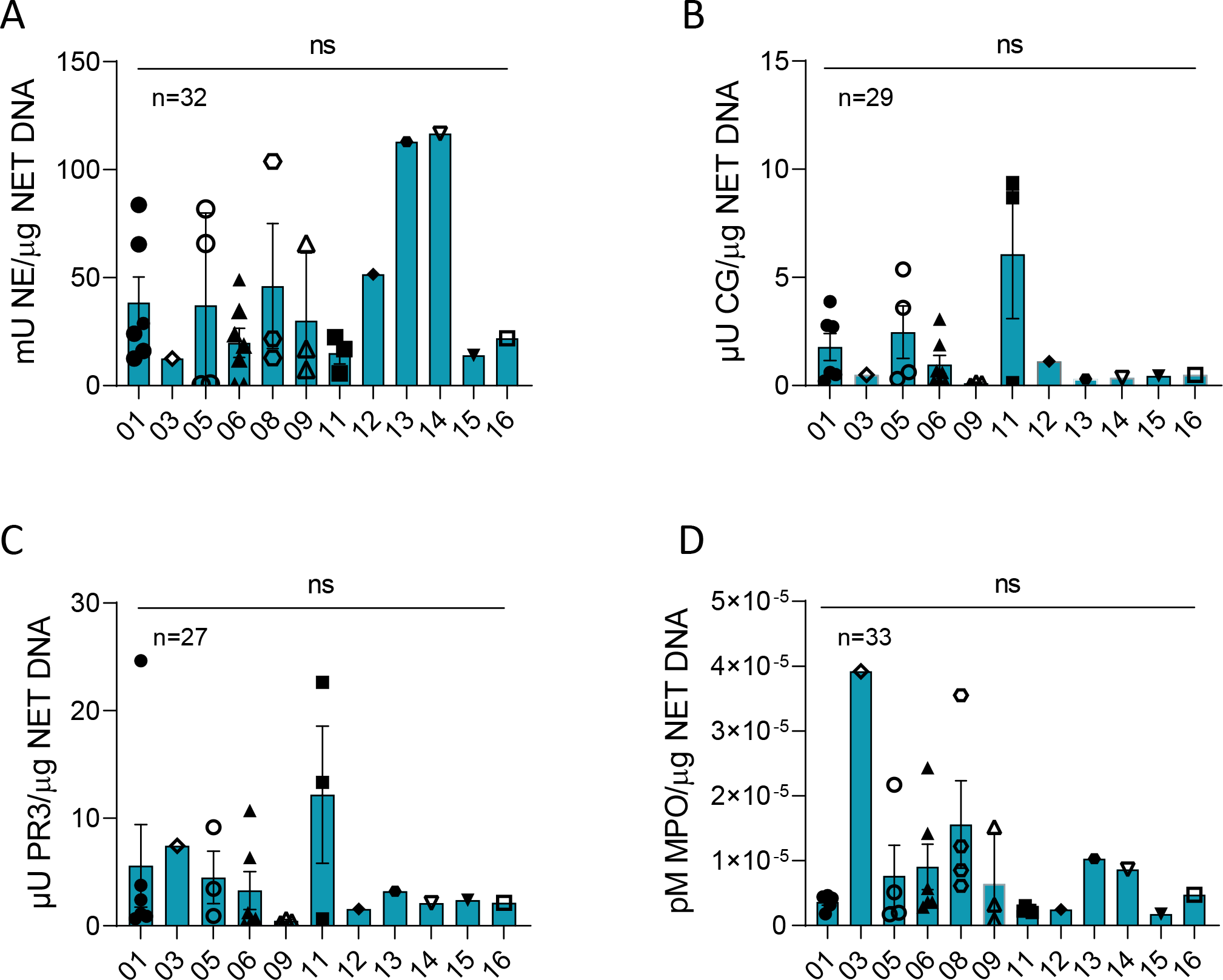
NET enzyme a*ctivity varied within but not across human donors*. **A)** Human NE (p=0.129), **B)** CG (p=0.188), **C)** PR3 (p=0.718), and **D)** MPO (p=0.143) enzyme activities were measured in NETs isolated from up to 12 normal donors between 27 to 33 blood draws and normalized per μg DNA and were not significantly different between donors. NE data analyzed across subjects by one-way ANOVA. PR3, CG, and MPO data analyzed across subjects by Kruskal Wallis.

**Figure 4.**
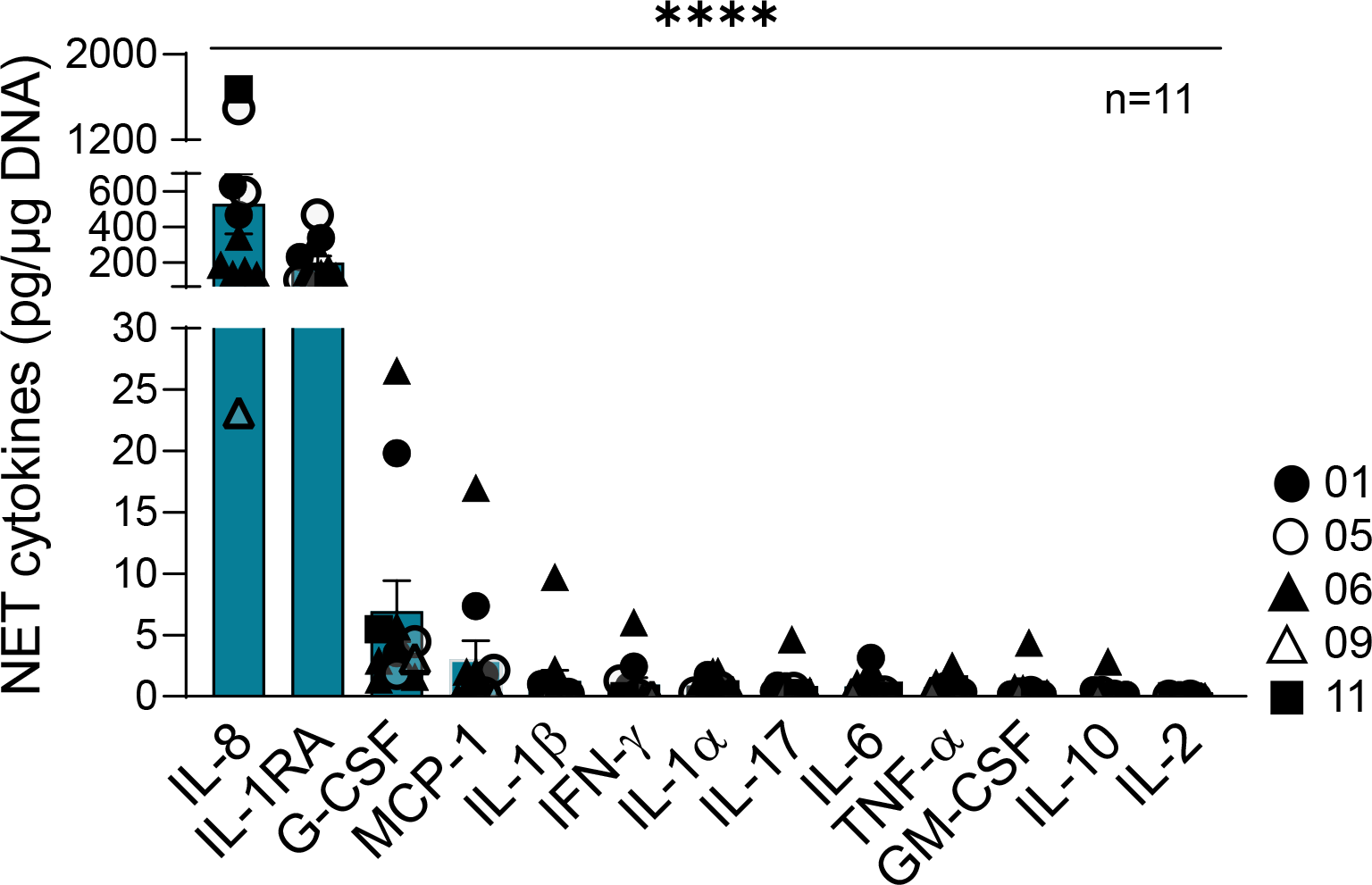
IL-8, IL-1RA and G-CSF were consistently present across all donor NETs. The concentration of cytokines was measured in isolated NETs from 5 donors across 11 blood draws and were normalized per μg DNA (p=<0.0001). Data analyzed across subjects by one-way ANOVA.

**Figure 5.**
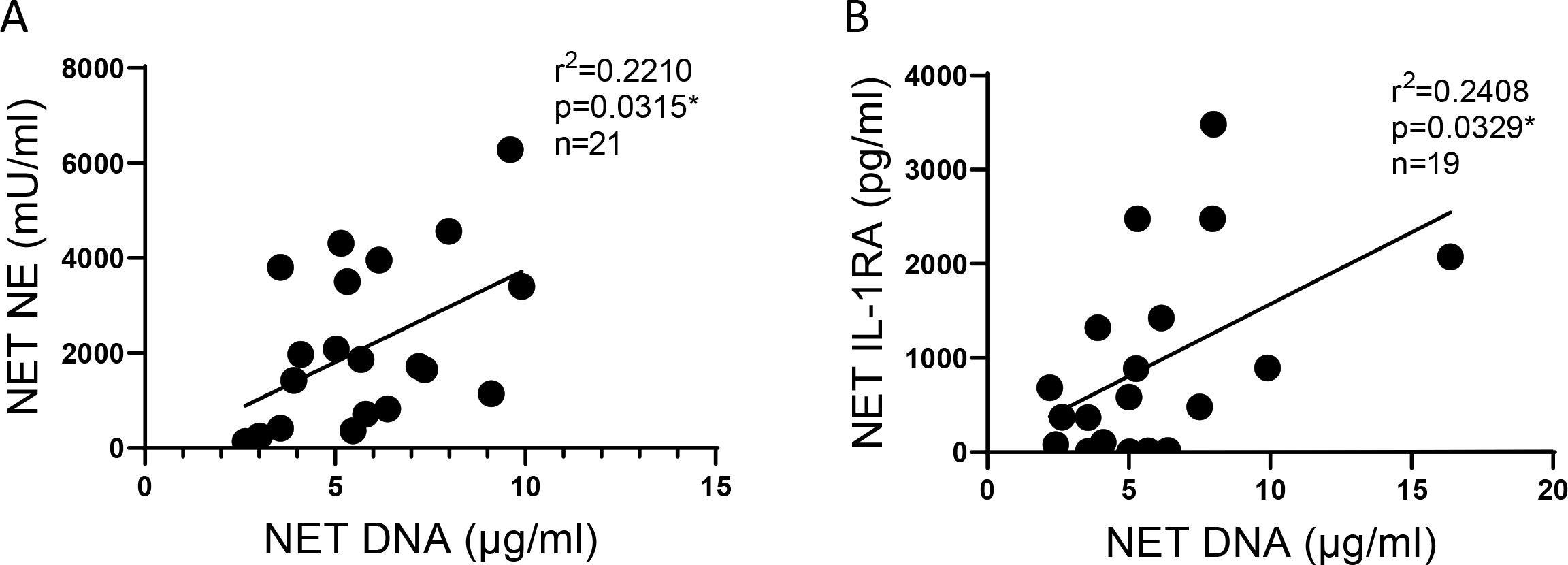
NE and IL-1RA correlate with NET DNA concentration. **A)** NE activity correlates with DNA concentrations in NETs from 6 blood donors across 21 blood draws. **B)** The anti-inflammatory cytokine IL-1RA concentration correlates with DNA concentrations in NETs from 7 blood donors across 19 blood draws. Data analyzed by Pearson correlation with best fit line by linear regression.

NETs contribute cytokines to the extracellular milieu. In Figure 4, we report the cytokines present in NETs, normalized to μg of DNA. The most abundant cytokines in isolated NETs were IL-8, a potent neutrophil chemoattractant, and IL-1RA, a competitive antagonist for the IL-1 receptor 1 (12, 20). Isolated NETs from different donors all contained small amounts of G-CSF (1-26pg/ml), which is a potent stimulator for the bone marrow to increase neutrophil production (21). NET-bound cytokines could contribute to a positive feedback loop of inflammation. We found that increasing IL-RA concentrations correlated with increasing NET DNA concentrations (Figure 5B). We detected little or no GM-CSF, IFN-γ, IL-1*α*, IL-1β, IL-2, IL-6, IL-10, IL-17A, MCP-1, or TNF*α* proteins in NETs from healthy donors.

### 2.4. NETs from different human subjects have varying susceptibilities to serine protease inhibitors

NETs isolated from the peripheral blood of healthy donors were assessed for susceptibility to protease inhibitors, or serpins (Figure 6A-C). Alpha-1 antitrypsin (AAT) is a naturally occurring, irreversible protease pseudosubstrate that binds and inactivates serine proteases. AAT is also known to inhibit apoptosis, and bind and inactivate IL-8 (22, 23). CG inhibitor (CGI) is a selective and reversible inhibitor of CG, having little or no effect on related proteases (24). NE inhibitor (NEI) is an N-benzoylindazole derivative that selectively targets the binding domain of NE (25). Sivelestat (SIV) is a selective reversible inhibitor of NE (26).

**Figure 6.**
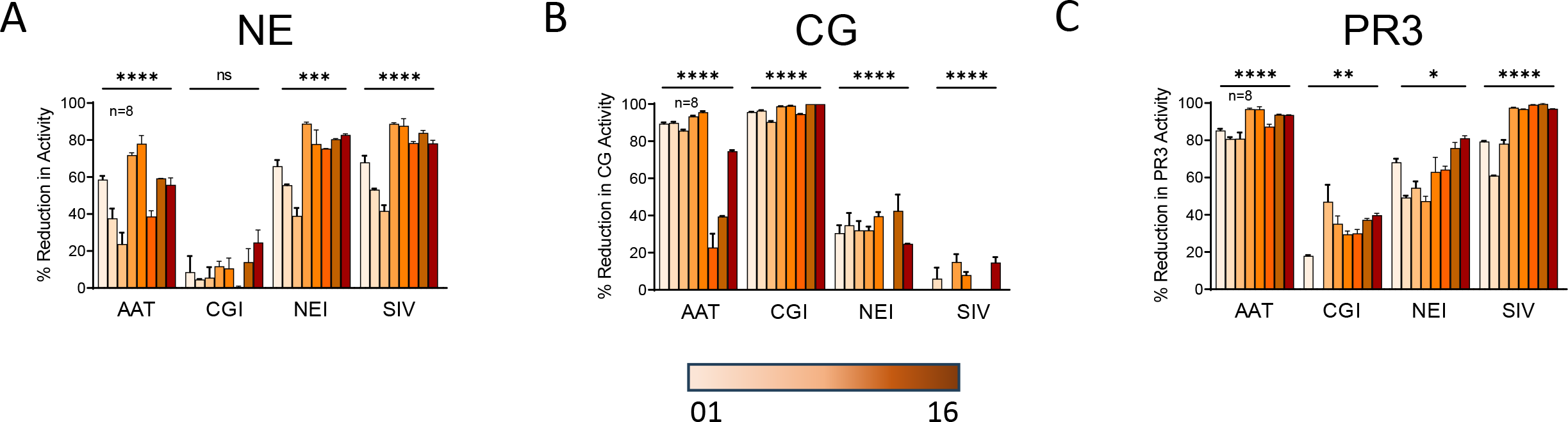
NETs from different subjects vary significantly in their susceptibility to protease inhibitors. In cell-free assays, 100μg/ml protease inhibitors AAT, CGI, NEI, and SIV were incubated 18 hours with NETs isolated from 8 donors. Samples were then assayed for **A)** NE, **B)** CG, and **C)** PR3 activities and are expressed as percent reduction in enzymatic activity. Data analyzed across subjects by one-way ANOVA *p<0.05, **p<0.01, ***p<0.001 ****p<0.0001.

The mean reductions in NE, CG and PR3 activities were different across donors for most inhibitors (Figure 6A-C). We observed that some “selective inhibitors” cross-reacted and inhibited other serine proteases. For instance, CGI exposure, which was highly effective in blocking CG activity in all donor NETs, also reduced PR3 activity, to a lesser degree (Figure 6B). Notably, although we anticipated that AAT would decrease all serine proteases substantially, AAT had the greatest impact on PR3 activity (Figure 6C), less effect on CG (Figure 6B), and the smallest impact on NE (Figure 6A).

### 2.5. NET components and correlation to barrier dysfunction

We have previously reported that HBE exposed to NETs have a significant decrease in barrier function as measured by the TEER of epithelial monolayers (10). However, it was unknown if NET components across multiple donors would demonstrate similar associations with loss of epithelial barrier function. We therefore sought to correlate which NET components from multiple donors were associated with reduced TEER. Increasing DNA concentrations in NETs negatively correlated with TEER in HBE (Figure 7A) however, activity of NE, the predominant protease of NETs, did not correlate with reduced TEER (Figure 7B). While the considerable variability in NE activity across NET donors impacted this correlation, it is unlikely that a single component of the complex NET structure would predict monolayer resistance. This suggests that cell junction damage from neutrophil NETs is dependent on multiple bioactive molecules. In addition, IL-1RA in NETs trended towards correlating positively with TEER, which suggests IL-1RA in the NETs may offer a protective effect to the bronchial epithelia (Figure 7C). Concentrations of CG, PR3, MPO, and LDH in NETs did not correlate with reductions in TEER (data not shown)

**Figure 7.**
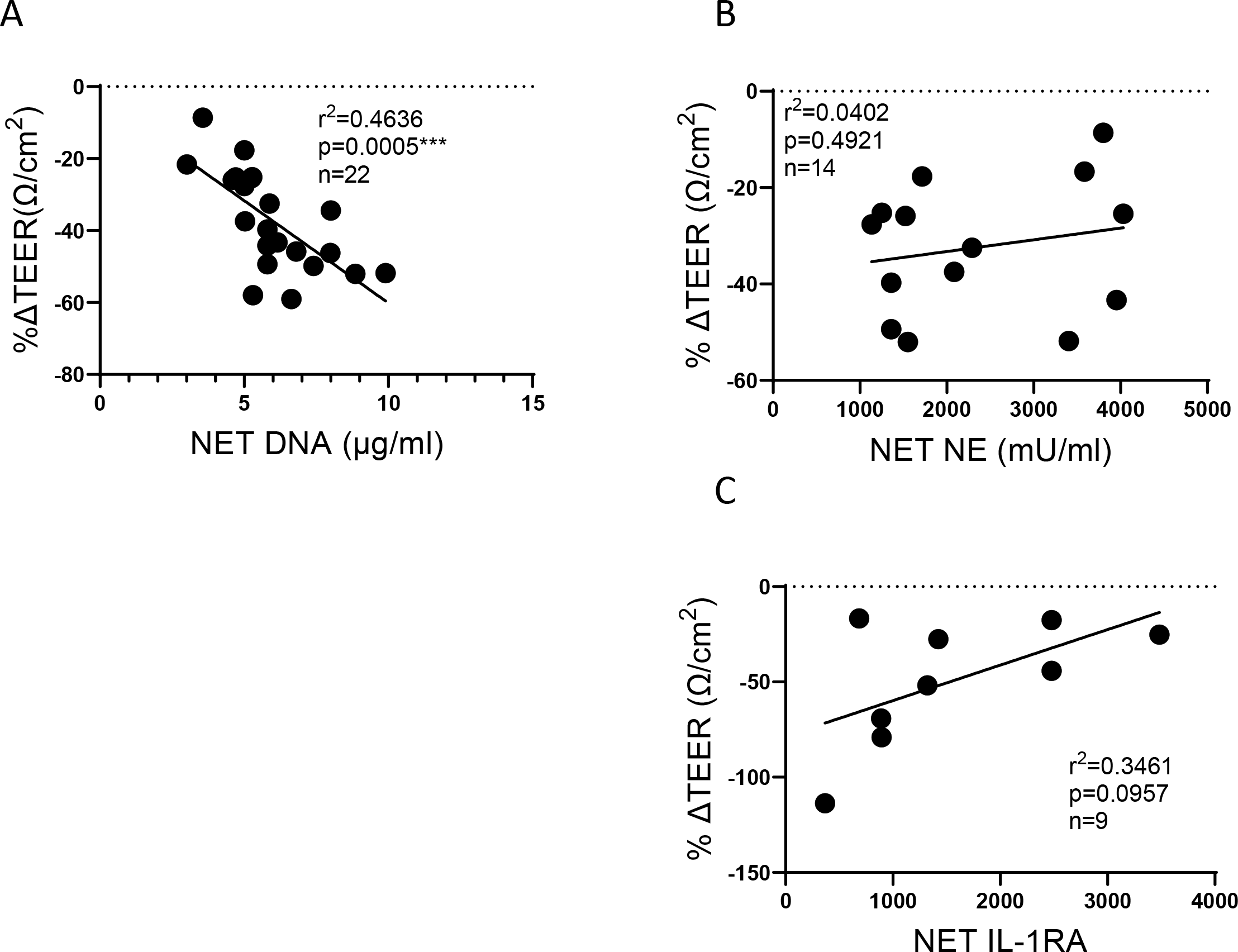
Correlation of NET components to change in lung epithelial barrier function. HBE grown at air-liquid interface (ALI) and exposed to PBS control or 5μg/ml NETs from up to 7 donors across 9 to 22 blood draws. **A)** The DNA concentration in NETs exposed to HBE significantly correlated to a reduction in TEER (r^2^=0.4536, p=0.0005). **B)** Activity of NE, the predominant serine protease in NETs, did not correlate with barrier function. **C)** Increasing NET concentrations of the anti-inflammatory cytokine IL-1RA trended toward preservation of TEER. Data analyzed by Pearson correlation with best fit line by simple linear regression.

## 3. Discussion

Elevated NETs are present in many lung diseases including Cystic Fibrosis (CF), non-CF bronchiectasis, asthma and COPD (27-30). We and others have demonstrated that NETs can disrupt barrier function of the epithelium and endothelium, a key mechanism of NET-induced pathogenesis in the lung (10, 31). NETs are complex structures with over 100 proteins, many of which are immunomodulatory, so defining NET concentrations is challenging. Herein, we describe the spectrum of NET contents for healthy human subjects and determine the critical components most associated with disrupted lung epithelial barrier function.

NETs contain double and single-stranded (ss) genomic DNA, which may have differential impacts on the host. We found that dsDNA was approximately one third of the total DNA present in the NETs of healthy donors. Susceptibility of ssDNA or dsDNA to host endonucleases varies considerably and ssDNA is often less stable. This may translate to NETs with greater dsDNA content having longer half-lives in vivo, amplifying their biologic effect (32-34). ssDNA and dsDNA can be perceived by the body differently and could elicit varying immune responses. “Uptake” of exogenous DNA by bacteria, e.g. Pseudomonas, can be beneficial as DNA provide key nutrients for growth (35). It is unclear if bacteria can more easily uptake ssDNA vs dsDNA, but the NET ssDNA vs dsDNA content could alter bacterial pathogenesis.

LDH levels were high in the NETs of our human subjects. The amount of LDH we reported is very consistent across our healthy donors. The NET concentration of LDH may also be related to the stimulus for NETosis, which in our case is PMA. Although PMA can cause necrosis of neutrophils, at the low doses used in this study no necrosis was observed by western blot. LDH is widely used as a measure of cytotoxicity, but cytotoxicity may be overestimated in experiments if the NET contribution is not accounted for in interpretation of results (36).

There were considerable differences in variability in NET enzymatic activity within the same person from day-to-day. The factors that regulate the extent of enzymatic activity within NETs on different days are unknown. We investigated susceptibility to protease inhibitors because we postulated that steric hindrance of the NET structure may limit direct binding and protease inactivation. We found significant differences in effectiveness of protease inhibitors, which did not trend with the magnitude of protease activity. Interestingly, NE activity had the most variable response across donors to both irreversible (AAT) and reversible (SIV) inhibitors. NE has been shown to be the most abundant serine protease in NETs and in this study we demonstrate that NE is the greatest contributor of serine protease activity found in healthy subjects’ NETs. The reduction of NE activity by the inhibitors we tested was highly variable across subjects. The difference in variability across donors to NE inhibitors could have a considerable impact in the utility of anti-protease agents as therapies to reduce NET-induced pathology in CF or AAT deficiency. Further work will be needed to better understand what affects NE susceptibility to various protease inhibitors.

It would be impractical to account for the activity of all the components of NETs. This makes determining a given NET “dose” challenging; thus, many researchers use DNA content to define “NET dose.” We found that the NET NE and NET IL-1RA correlated with DNA content. The DNA concentration best predicted disruption of the barrier function of HBE, providing further evidence for using DNA to determine NET concentrations. An alternative approach is to pool the NETs from multiple donors and thus average the protein activity differences.

Cytokine content in NETs was consistent across five donors (eleven NET isolations). IL-8 was the most abundant cytokine and is a critical neutrophil chemoattractant known to be elevated in many inflammatory lung diseases including CF and acute respiratory distress syndrome (ARDS) (37-39). *In vivo*, IL-8 can be free or bound, e.g. to immunoglobulin or heparin, which may affect the half-life and biologic impact (40, 41). It is unclear if IL-8 bound to DNA has similar caveats; and the extent to which a negatively charged DNA molecule could affect IL-8 interactions with the IL-8 receptor is unknown. Moreover, it is possible that the pool of IL-8 bound to DNA may not be detected to the same magnitude depending on the epitope by which IL-8 is detected for a given assay, thereby underestimating the IL-8 concentration of a given compartment. Another proinflammatory cytokine we found in the NETs of our donors was G-CSF, which to our knowledge, has not been previously reported. G-CSF is critical for neutrophil maturation and stimulating bone marrow to generate and release neutrophils. Both G-CSF and IL-8 can decrease neutrophil apoptosis, extending the neutrophil lifespan (42, 43). NETs also contained a considerable concentration of the anti-inflammatory cytokine IL-1RA. IL-1RA binds to the IL-1 receptor I, but does not activate signal transduction as the IL-1*α*/β agonists do. We observed a correlation between higher NET IL-1RA and reduced disruption of epithelial barrier function. It is possible that the IL-1RA in the NET may directly limit NET-induced barrier dysfunction by decreasing NET driven IL-1*α*/β signaling. We previously demonstrated that IL-1*α*/β signaling drives epithelial secretion of TNF-*α*, which is known to alter epithelial tight junctions critical to barrier function (12, 44).

A strength of our study is the inclusion of donors across the age spectrum and of both genders. Another strength was a study period of over 3-years with multiple draws from twelve donors. All donations were at least 1 month apart hoping to minimize the impact of any single event on our results. We also took every effort to limit known factors that affect neutrophil function within our population, including collecting all blood donations within the same 2 hour window in the morning. We recruited nonsmokers, with no consumption of alcohol or NSAIDS within 3 days of donation. An equal number and average age of male and female donors (6M average age of 41, and 6F average age of 38) were used with a stipulation of greater than 2 weeks after an acute illness or receiving antimicrobial therapy (45-47). The limitations of our study include the inability to control all the factors that influence an individual’s neutrophil function including subclinical infections, which could explain some of the variation seen. Despite the fact that we chose a healthy population with no major medical problems, we could not limit all prescribed medications nor could we control for stress or other inflammatory stimuli. Another limitation is the limited sample size. For example, our study did not find a significant heterogeneity in neutrophil numbers; however, given the variance estimates, we would expect to have 80% power to detect a significant heterogeneity in NET total DNA content if we have 12 subjects each with 8 repeated measurements. In this regard, our study generates important preliminary data to inform future studies related to NET biology and impact on epithelium. We acknowledge that the baseline phenotype of the donor neutrophil (including cellular maturity, activation, degranulation potential, etc.) could significantly alter the composition and concentration of NETs generated. Our fu-ture studies will focus on uncovering the degree to which these factors regulate observed differences in NETs.

In conclusion, we demonstrate that DNA content and susceptibility to protease inhibitors are significantly different across healthy donors. These findings are clinically relevant as DNAse and anti-proteases (such as AAT) are commonly used therapies to treat lung diseases including CF and AAT deficiency, respectively. Future studies will elucidate if variable responses to these therapies could be due to the heterogeneity of NET contents across human subjects that we report in this study.

## 4. Materials and Methods

### 4.1. Isolation of human neutrophils and NETs

The study protocol, no. 2016-3837, was approved by the University of Cincinnati institutional review board (IRB). Human peripheral blood was collected from healthy adult donors in sodium citrate vacuum tubes (Fisher Scientific). Neutrophils were isolated by negative magnetic bead selection using MACSxpress Neutrophil Isolation Kit, Human (Miltenyi). Cell counts and viability were assessed using Trypan blue, 0.4% (Fisher Scientific). PMNs were suspended in RPMI-1640 media (GIBCO) with 3% fetal bovine serum (Fisher Scientific) at a concentration of 5x10^6^ cells/ml. Cells were incubated at 37°C, 5% CO_2_, 4h in 100mm tissue culture-treated petri dishes (Fisher Scientific) with 25nM PMA (Sigma) added to stimulate NETosis (12, 13). Following incubation, the viscous NET layer was washed twice with PBS, scraped from the dish, and mixed vigorously. The sample was then centrifuged at 450xg, 10m, 22°C to remove cell debris. Cell-free NETs suspended in PBS were added to the apical surface of HBE.

### 4.2. Primary epithelial cell culture

Primary HBE were provided by the CCHMC CF Research Development Program Translational and Model Systems Cores. Cells were cultured as previously described (14). Primary HBE from normal donors were obtained from University of North Carolina Airway Cell Core and were utilized at passages 2 to 3. HBE were differentiated and grown at ALI on 6.5mm Transwell inserts (Costar). The apical surfaces of HBE were exposed to PBS or NETs suspended in PBS at 37°C, 5% CO_2_ for 18 hours unless otherwise noted. Supernatants and cell protein lysate were stored for future testing. The use of primary HBE was approved by the CCHMC Pulmonary Biorepository Core, and University of Cincinnati IRB.

### 4.3. Transepithelial electrical resistance

TEER was measured across each epithelial cell layer in duplicate before and after exposure to test conditions and PBS control using an Epithelial Volt/Ohm meter EVOM2 and chopsticks electrode set STX3 (World Precision Instruments; Brewington 2018). Results are expressed as percent change from baseline TEER values.

### 4.4. Characterization of NETs and cell supernatants

Samples of NET isolates and HBE supernatants were collected and aliquots stored at -80°C. HBE were processed for protein lysates and stored at -20°C (Complete Lysis-M Roche diagnostics). NET dosage was an average of extracellular dsDNA concentration (QuantiFluor ONE dsDNA System, Promega) and total DNA content (SYTOX green nucleic acid stain, Fisher Scientific, (15). Protein content of NETs and cell lysates was determined using DC Protein assay (BioRad).

NETs were assayed for: LDH (Promega CytoTox 96 Cytotoxicity Assay); MPO activity (Cayman Chemical MPO Activity Assay); NE, PR3, and CG activities (fluorescent resonance energy transfer (FRET) substrates Abz-APEEI MRRQ-EDDnp (NE), AbzVADnV RDRQ-EDDnp (PR3), Abz-EPF WEDQ-EDDnp (CG) Peptide Institute; (16). Cytokine analysis was performed by Cincinnati Children’s Hospital Research Flow Cytometry Core per kit instructions (Milliplex MAP human protein panel, Millipore-Sigma).

### 4.5. Microscopy

Neutrophil and NET images were captured on a Zeiss Axioplan 2 or a Nikon A1R GaAsP inverted SP microscope using antibodies to NE (Millipore 481001), MPO (Abcam 25989) with AF568 and AF488 secondary antibodies (Invitrogen) and Hoechst nucleic acid stain (Invitrogen).

### 4.6. Statistical analysis

Statistical analysis was performed using GraphPad Prism software v7.04 and SAS version 9.4 (SAS Institute, Inc, Cary, NC). Subject means were compared using ordinary one-way ANOVA with Bonferroni corrections for multiple comparisons.(48) We assessed model assumptions using residual plots in the ANOVA models, and used Kruskal-Wallis tests for outcomes that did not meet the normality assumption (49).Variability between subjects’ values was measured by comparing standard deviations using the Brown-Forsythe test. Correlations were analyzed using Pearson correlations with best-fit line by simple linear regression. P-values ≥0.05 were not considered significant, while significant p-values are denoted by: <0.05*, ≤0.01**, ≤0.001***, ≤0.0001****. Graphs include data from at least three separate experiments unless indicated. Each donor is represented by the same symbol across all graphs.

### 4.7. Research Compounds

AAT (UniProt# P01009, Cayman Chemical), NEI (CAS#1448314-31-5, Cayman Chemical), SIV (CAS# 201677-61-4, Cayman Chemical), CGI (CAS# 429676-93-7, Cayman Chemical), PMA (CAS #16561-29-8, Millipore-Sigma).

### 4.8 Study Population

Healthy male and female adult subjects were enrolled in our study from January 2020-August 2023 who met the following criteria: healthy, nonsmokers, no history of malignancy, no history of autoimmune disorders, no use of immunosuppressive agents (including steroids) within 90 days, no acute illness or antimicrobials within 2 weeks. Subjects had to abstain from alcohol or NSAIDS for 72 hours prior to donation.

## Supplementary Materials

The following supporting information can be downloaded at: www.mdpi.com/xxx/s1, Table S1: Characteristics of Healthy Human NET donors.; Table S2: Quarterly blood collection calendar.; Figure S1: Purified neutrophils prior to stimulation to NETosis.

## Author Contributions

Designing research studies, KH, MC, and MI. Conducting Experiments, MC and MI. Analyzing data, MC, MI, EKo, and KH. Intellectual content, MC, MI, EKr, JH and KH. Statistical analyses, NZ. Writing the manuscript, MC, MI and KH. Revisions, all authors. All authors contributed to the article and approved the submitted version.

## Institutional Review Board Statement

The study protocol, no. 2016-3837, was approved by the University of Cincinnati IRB. Healthy adult human subjects provided written informed consent prior to participation in this study.

## Informed Consent Statement

Subjects provided informed consent to participate in this study.

## Data Availability Statement

**The data presented in this study are available on request from the corresponding author.**

## Acknowledgments

We would like to thank Jessica Meeker, Hunter Morgan, Alicia Ostmann and Dr. John Brewington from the CCHMC CF Research Development Program Translational and model Systems Core for culturing and providing HBE cells and Alyssa Sproles from Cincinnati Children’s Research Flow Cytometry Core.

## Disclaimer/Publisher’s Note

The statements, opinions and data contained in all publications are solely those of the individual author(s) and contributor(s) and not of MDPI and/or the editor(s). MDPI and/or the editor(s) disclaim responsibility for any injury to people or property resulting from any ideas, methods, instructions or products referred to in the content.

---

**Supplemental Table 1.** Characteristics of Healthy Human NET donors. Peripheral blood was freshly drawn for immediate isolation of neutrophils with subsequent NET generation from 12 healthy male and female donors. Donor ages ranged from 21-64.

| **Donor** | **n** | **Gender** | **Age** |
| --- | --- | --- | --- |
| **01** | 5 | F | 49 |
| **03** | 4 | F | 43 |
| **05** | 4 | M | 52 |
| **06** | 6 | F | 63 |
| **08** | 4 | M | 34 |
| **09** | 3 | M | 24 |
| **11** | 4 | M | 21 |
| **12** | 1 | M | 64 |
| **13** | 1 | M | 52 |
| **14** | 1 | F | 24 |
| **15** | 1 | F | 25 |
| **16** | 1 | F | 23 |

**Supplemental Table 2.**
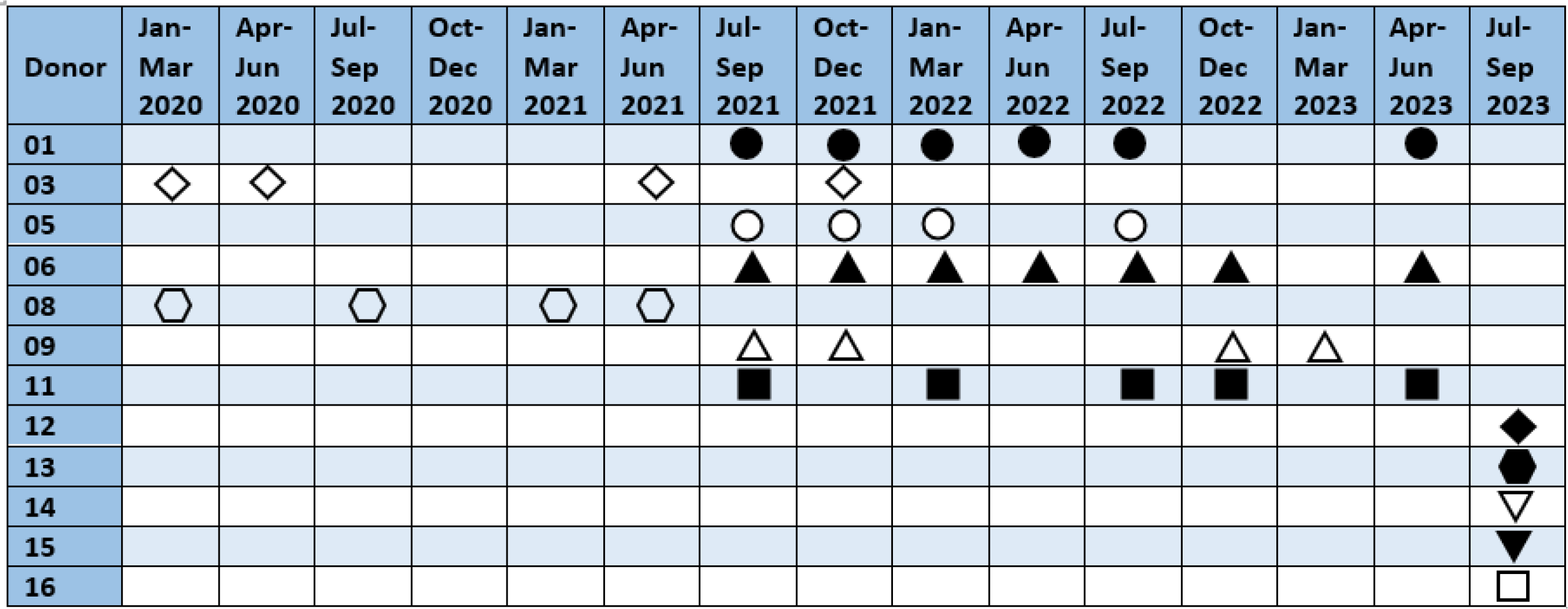
Quarterly blood collection calendar. Time intervals of peripheral blood collection from enrolled study subjects between 2020-2023.

**Supplemental Figure 1.**
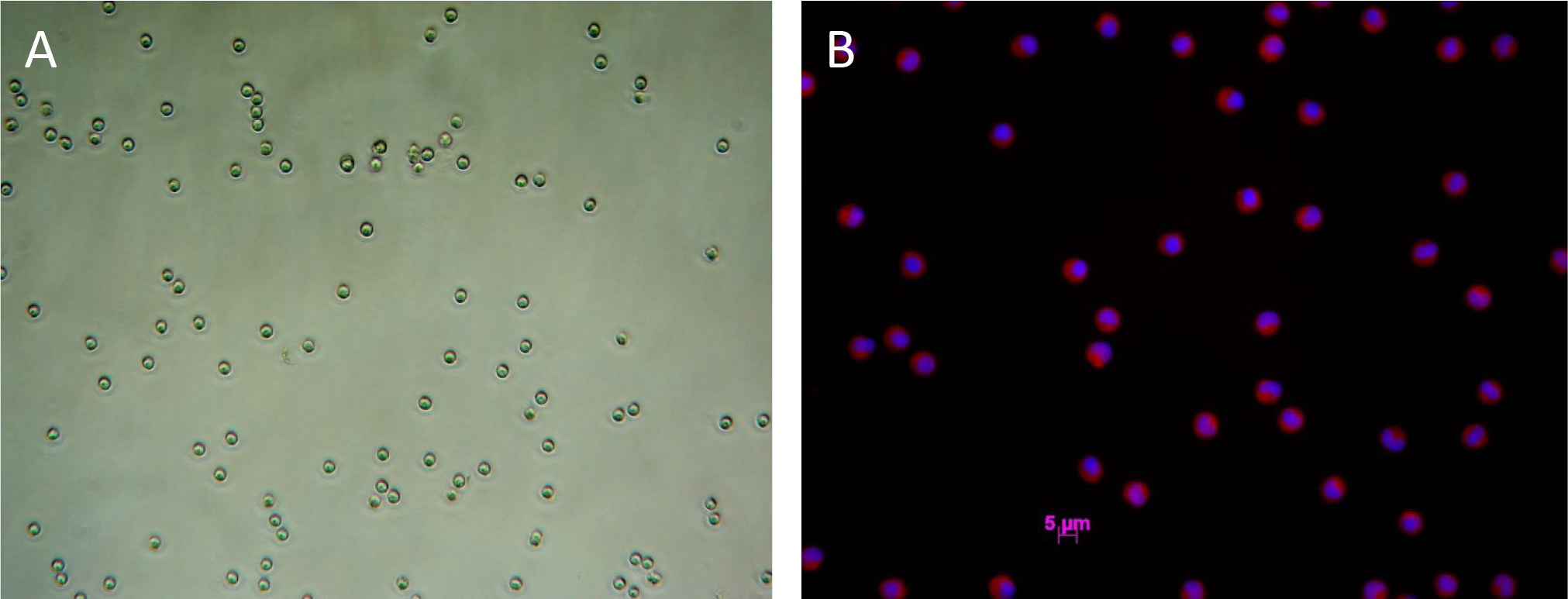
Purified neutrophils prior to stimulation to NETosis. **A)** Bright field image of enriched unstained neutrophils obtained from donor blood using negative magnetic bead selection and suspended in PBS with 0.04% trypan blue. **B)** Merged immunofluorescent image of unstimulated neutrophils at a concentration of 5x10^6^/ml and a density of 1x10^5^/cm^2^ stained with NE antibody (AF568, red), and Hoechst (blue).

